# Layered stomatal immunity contributes to resistance of *Vitis riparia* against downy mildew *Plasmopara viticola*

**DOI:** 10.1101/2025.02.23.639706

**Authors:** Wei Ji, Wei Zheng, JunJie Mei, Xiaoyu Liu, Naomi Abe-Kanoh, Mohammad Saidur Rhaman, Guochen Qin, Wenxiu Ye

## Abstract

Downy mildew (DM), caused by *Plasmopara viticola*, is one of the most serious grapevine diseases. Resistant grapevines are a well-known tool for mitigating pathogen-caused damage. We evaluated 29 global grapevine cultivars from 7 species for the sensitivity to *P. viticola*. Chardonnay belonging to the sensitive species *V. vinifera* and Qingdahean belonging to the well-known resistant species *V. riparia* were chosen for further investigation on the resistance mechanism against DM. Unlike Chardonnay, Qingdahean exerted an inhibitory effect on stomatal targeting, suppression of stomatal closure, stomatal penetration of *P. viticola*, and the development of primary hyphae and haustoria during the early phase of infection, and contained higher levels of malondialdehyde (MDA), which was significantly increased by *P. viticola* infection, toxic to the pathogen and had an interfering effect on the stomatal targeting. Furthermore, Qingdahean resisted pathogen invasion through the rapid induction of guard cell death and the hypersensitive responses (HR) of other cell types. These findings suggest that resistance to *P. viticola* consists of layered stomatal immunity in addition to the well-known HR in *V. riparia*, which is overcome by the pathogen in *V. vinifera*.

## INTRODUCTION

The grapevine (*Vitis*) is an economically important fruit tree cultivated worldwide. It yields an impressive yearly production of 77 million tons of grapes, covering a total area of 7 million hectares (FAO, 2021). Grapevine downy mildew (DM) is one of the most devastating and economically alarming diseases in the grape industry, which is caused by *Plasmopara viticola*, a biotrophic oomycete, resulting in catastrophic yield losses and significant defects in berry quality (Peng et al. 2023; Guo et al. 2024). During the grapevine growing season, asexual propagation of *P. viticola* is responsible for the efficient spread of *P. viticola*. At high humidity and warm temperatures, the lemon-shaped sporangia (asexual reproduction structure) release flagellate zoospores that swarm and specifically target stomata for penetration. Once zoospores reach stomata, they shed their flagella, form cysts, and grow germ tubes into the substomatal cavity. The germ tubes then dilate into infection vesicles, from which primary hyphae emerge, spread within the leaf tissue, primarily in the intercellular spaces of the spongy parenchyma, and form haustoria, taking nutrients from host cells. After a few days, sporangiophores protrude from the stomata and form sporangia, and a new secondary infection occurs (Burruano 2000; Gessler et al. 2011; Armijo et al. 2016; Yin et al. 2017).

Stomata, formed by a pair of guard cells, are main invasion point of infection for many pathogens causing important diseases such as many bacterial diseases, wheat rust and grapevine downy mildew (Kiefer et al. 2002; Melotto et al. 2006; Ye et al. 2020). A known defense strategy is that guard cells recognize the so-called microbe-associated molecule patterns (MAMPs) from bacteria and fungi, such as the bacterial flagellin peptide, fungal cell wall component chitin and chitosan, and induce stomatal closure and guard cell death to ward off pathogen infection (Melotto et al. 2006; Ye et al. 2020). To overcome stomatal closure, it is shown that bacteria and fungi secret toxins (e.g. fusicoccin, coronatine, and syringolin A) and effectors (e.g. AvrB, HopF2, HopX1, and HopZ1) to reopen stomata for penetration (Hou et al. 2024; Melotto et al. 2024). In the case of bacterial infection, stomatal reopening at the later infection stage is shown to be a defense strategy to disrupt the formation of an aqueous apoplast suitable for bacterium proliferation, while effectors (e.g. AvrE, HopM1) are secreted by the bacteria to recreate aqueous apoplast for proliferation (Zhang et al. 2019; Hu et al. 2022; Liu et al. 2022; Roussin-Léveillée et al. 2022). On the other hand, the form of stomatal immunity against pathogenic oomycete invasion is poorly understood.

Post-stomatal defenses involve the accumulation of toxic compounds against pathogens and signaling molecules to trigger various defense responses, including the hypersensitive response (HR), a form of regulated host cell death at the site of infection (Bi and Zhou 2021; Jones et al. 2024). In the incompatible interaction between grapevine and *P. viticola*, HR has been widely observed and is thought to limit the growth of the biotrophic pathogen (Ma et al. 2020; Fu et al. 2023; Paineau et al. 2023). Signaling molecules such as reactive oxygen species (ROS), malondialdehyde (MDA), and phytohormones, such as abscisic acid (ABA) and salicylic acid (SA), play important roles in both stomatal and post-stomatal defenses (Yu et al. 2016). It remains poorly understood how these molecules are involved in the grapevine resistance to DM.

In the present study, we explored resistance characteristics of 29 grapevine germplasms and investigated the resistance mechanism of *V. riparia* against DM by comparing with the susceptible species *V. vinifera*. The findings indicate that, apart from HR, stomatal immunity, which consists of at least four separate layers, has a role in the resistance to *P. viticola* in *V. riparia*. This underscores the grapevine downy mildew disease as a valuable model for studying stomatal immunity against pathogenic oomycetes.

## RESULTS

### Selection of cultivar and their different responses to *P. viticola*

We collected 29 grapevine cultivars covering 7 species from various parts of the world including *V. vinifera, V. rotundifolia, V. berlandieri, V. rupetris, V. labrusca, V. riparia,* and *V. amurensis* and investigated their resistance to *P. viticola* (Figure S1A, S2, Table S1). The inoculation was performed according to previous study (Paineau et al. 2022; Guo et al. 2024). Each leaf disc (8 mm) was inoculated with 20 μL *P. viticola* suspension in water of 1 × 10^5^ sporangia/mL on the abaxial surface. At 10 days after inoculation (DAI), the symptom of each leaf discs was imaged and the number of sporangia was counted. The findings showed that different cultivars produced varying numbers of sporangia per leaf disc (Figure S1B, S3). The highest number of sporangia were found in the Chardonnay (*V. vinifera*) cultivar, while the lowest amount was found in the cultivars Fry, Granny Val, 110R, Beta, 1103P, 5BB, and Qingdahean (*V. riparia*). Since *V. riparia* is well known for its resistance to DM, we decided to further investigate its resistance mechanism by comparing to the most vulnerable cultivar, Chardonnay. Biochemical and physiological analysis indicated that Chardonnay and Qingdahean had similar responses in terms of leaf pigment, ETR, photosynthetic rate (A) and intercellular CO_2_ concentration (Ci) at different DAI of *P. viticola* (Figure S4, S5). On the other hand, the two grapevines responded differently regarding transpiration rate (E), stomatal conductance (Gs), Fv/Fm, Phips2 levels (Figure S4, S5), suggesting that these parameters are involved in the immunity of *V. riparia* to *P. viticola*. In addition, the stomatal density of Qingdahean is significantly higher than that of Chardonnay (Figure S6).

### Stomatal targeting of zoospores is inhibited in *V. riparia*

In order to investigate the early resistance to *P. viticola* infection, the stomata targeting of zoospores was studied by Blankophor staining of leaves at the early timings from 0.5 to 9 hours after inoculation (HAI) in both Qingdahean and Chardonnay (Figure 1). The blue fluorescence indicates the stained *P. viticola* zoospores and the autofluorescence of stomata (Figure 1A). The findings showed that the stomatal targeting rate on Qingdahean of 44.6% was significantly less efficient compared to that of 80.5% on Chardonnay at 0.5 HAI (Figure 1B). Over the period of infection, the stomatal targeting rate rose in both cultivars and reached a plateau after 4 HAI. Nevertheless, the stomata targeting rate in Qingdahean (80.3%) was significantly lower than that in Chardonnay (93.5%). Taken together, these results suggest that during the initial phase of *P. viticola* infection, the stomatal defenses of *V. riparia* impede the successful targeting of zoospores to the stomata.

**Figure 1.**
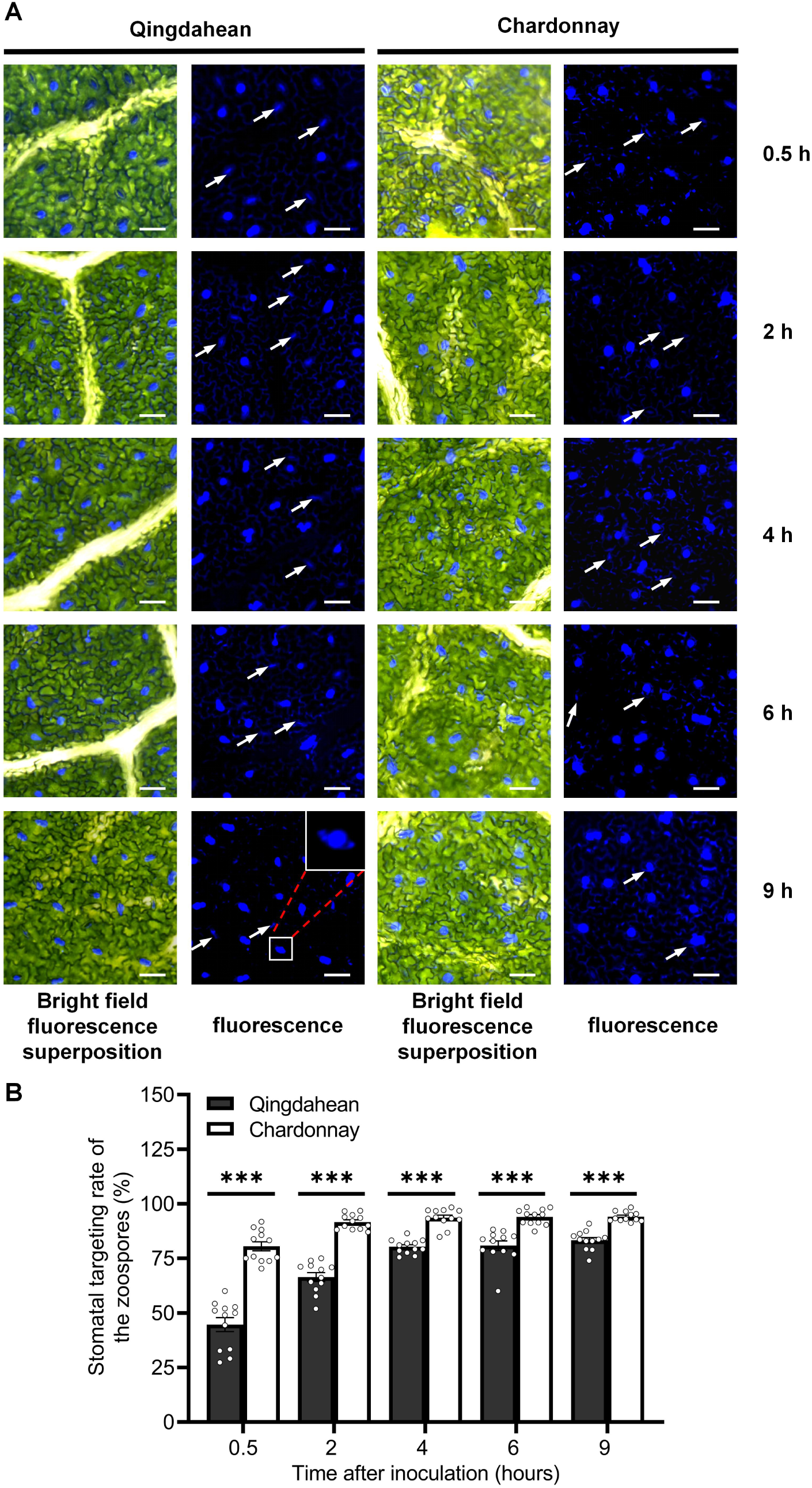
Stomatal targeting rate of *P. viticola* on Qingdahean and Chardonnay leaves. **(A)** Representative images of the encysted spores on the stomata at different hours post inoculation. Blue fluorescence indicates the stained spores by Blankophor. White arrows, autofluorescence of guard cells. Note that not all guard cells are indicated by arrows due to large number. Inset, an enlarged view showing zoospores localized on the stomata. Scale bar, 50 µm. **(B)** Quantification of the stomatal targeting rate of the *P. viticola* shown in A. Shown are individual data points and mean ± SE for n=12 over four independent leaf disks. The experiment was repeated three times with similar results (see Figure S13 for other independent sets of results). *\*\*\*P*<0.001, Student’s t-test.

### Suppression of stomatal closure by *P. viticola* is interrupted in *V. riparia*

Once the zoospores have successfully reached the stomata, they encyst and develop germ tubes penetrating into the stomatal cavity, where they generate substomatal vesicles followed by primary hyphae formation. Stomatal closure is an efficient strategy to ward off pathogen invasion, so we investigated the effect of *P. viticola* on stomatal aperture in Qingdahean and Chardonnay. As shown in Figure 2, the stomata without a germ tube closed rapidly upon inoculation in Qingdahean and Chardonnay. On the other hand, stomata targeted by germ tubes did not close up to 4 HAI in Chardonnay but closed rapidly after 0.5 HAI in Qingdahean. These results suggest that stomatal closure is an active defense strategy against *P. viticola* but is suppressed upon intimate physical interaction with the pathogen in *V. vinifera*. In contrast, *V. riparia* has a strategy to overcome the suppression.

**Figure 2.**
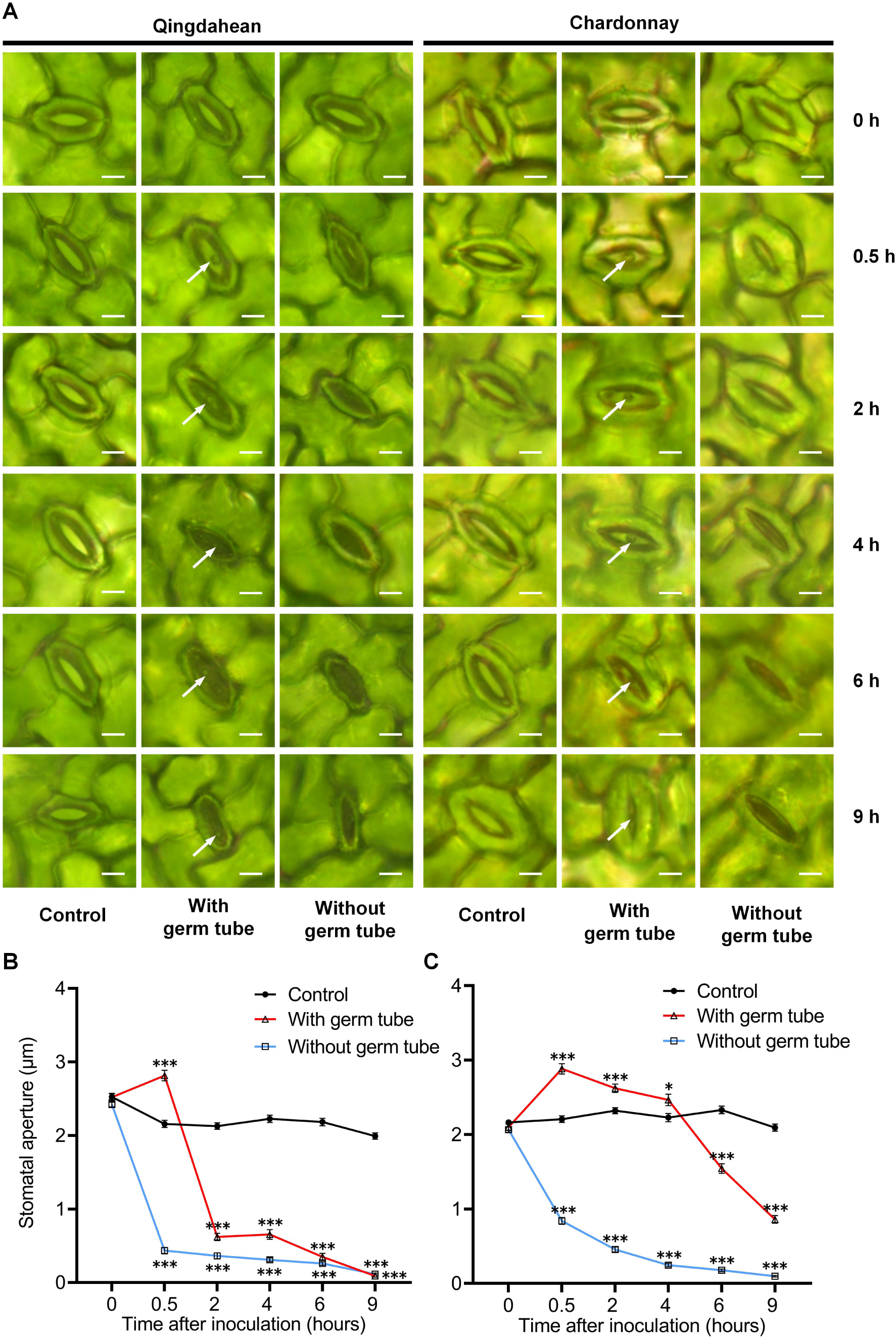
Stomatal movement induced by *P. viticola* on Qingdahean and Chardonnay leaves. Representative images of stomatal aperture at different hours post inoculation. White arrow, germ tube of *P. viticola*. Scale bar, 5 µm. **(B and C)** Quantification of the stomatal aperture of Qingdahean **(B)** and Chardonnay **(C)** shown in **A**. Shown are individual data points and mean ± SE for n=150 stomata over four independent leaf disks. The experiment was repeated three times with similar results (see Figure S13 for other independent sets of results). *\*P*<0.05, *\*\*P*<0.01, *\*\*\*P*<0.001, Student’s t-test compared with Control.

### Stomatal invasion of *P. viticola* and primary hyphae growth are suppressed in *V. riparia*

To determine if stomata can prevent the invasion of *P. viticola*, leaves were stained with trypan blue at different HAI, and the rate of stomatal invasion was then investigated by counting the rate of substomatal vesicle occurrence. It was shown that the rate of stomatal invasion in Qingdahean reached approximately 75% following successful stomatal targeting during a span of 2 HAI with *P. viticola* (Figure 3A, 3B). Subsequently, it further escalated to a plateau of 88.7% after 4 HAI. On the other hand, the rate of stomatal invasion in Chardonnay swiftly reached 96.2% within 0.5 hours, and swiftly reached 100% after 2 hours (Figure 3A, 3B).

**Figure 3.**
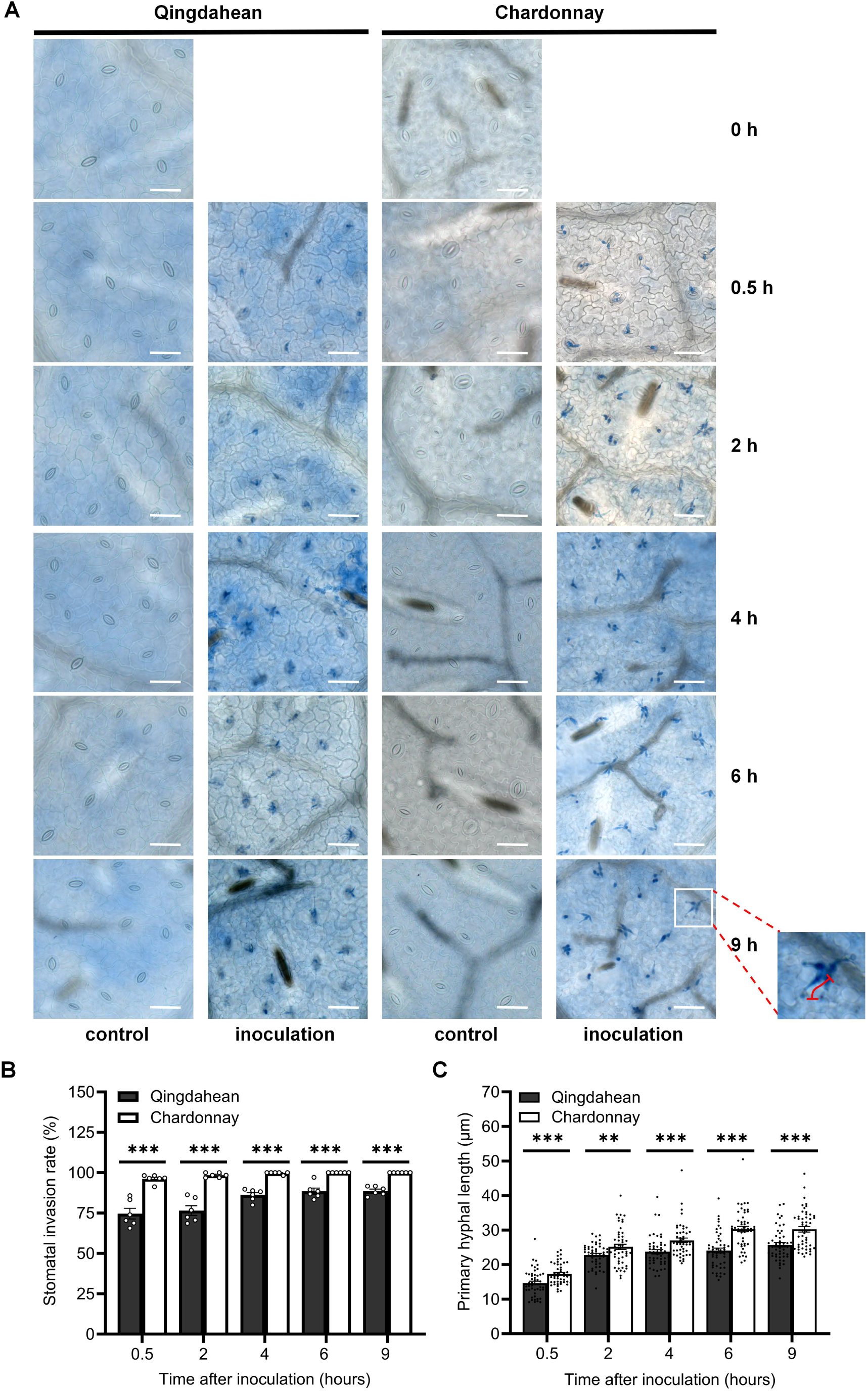
Stomatal invasion rate and primary hypha development of *P. viticola* in Qingdahean and Chardonnay leaves. **(A)** Representative images of *P. viticola* penetration through stomata at different hours post inoculation. Blue color indicates stained *P. viticola* tissues by trypan blue. Scale bar, 50 µm. The red line in the lower right corner illustrates the schematic for measuring the length of the primary hyphae. **(B)** Quantification of the stomatal invasion rate shown in A. Stomatal invasion rate was quantified as the ratio of the number of stomata having substomatal vesicles to the number of stomata targeted by zoospores. Shown are individual data points and mean ± SE for n=6 leaf disks. **(C)** Quantification of the length of primary hyphae shown in **A**. Shown are individual data points and mean ± SE for n=50 stomata targeted by zoospores over four independent leaf disks. The experiment was repeated three times with similar results (see Figure S13 for other independent sets of results). *\*\*P*<0.01, *\*\*\*P*<0.001, Student’s t-test. The experiment was repeated three times with similar results.

We further investigated whether the growth of primary hyphae is hindered in Qingdahean by measuring the length of primary hyphae. Over time, the primary hyphal length of Qingdahean and Chardonnay increased gradually (Figure 3A, 3C). However, the length of primary hyphae in Qingdahean was consistently shorter than that of Chardonnay. Taken together, these results suggest that the stomata of *V. riparia* act as a barrier against the invasion of *P. viticola* and the formation of primary hyphae to some degree.

### Guard cell death and post-stomatal HR contribute to the resistance of *V. riparia* to *P. viticola*

To further investigate the resistance mechanism of *V. riparia*, the morphological symptoms upon DM infection were observed. The results showed that the presence of obvious HR necrotic patches on the Qingdahean leaves occurred from 1 DAI, which were absent on healthy leaves (Figure 4A1-4B5, S8). The findings also indicated a progressive rise in both the size and quantity of necrotic lesions as the duration of the infection advanced. Moreover, the HR necrotic patches failed to generate sporangiophores and sporangia of *P. viticola*. In contrast, the Chardonnay leaves inoculated with *P. viticola* did not exhibit any necrotic patches. From 3 DAI, a few *P. viticola* sporangiophores were observed, and by 5 DAI, the entire leaves were completely covered by sporangiophores and sporangia (Figure 4). These findings indicate that HR contributes to the defense of *V. riparia* against *P. viticola*.

**Figure 4.**
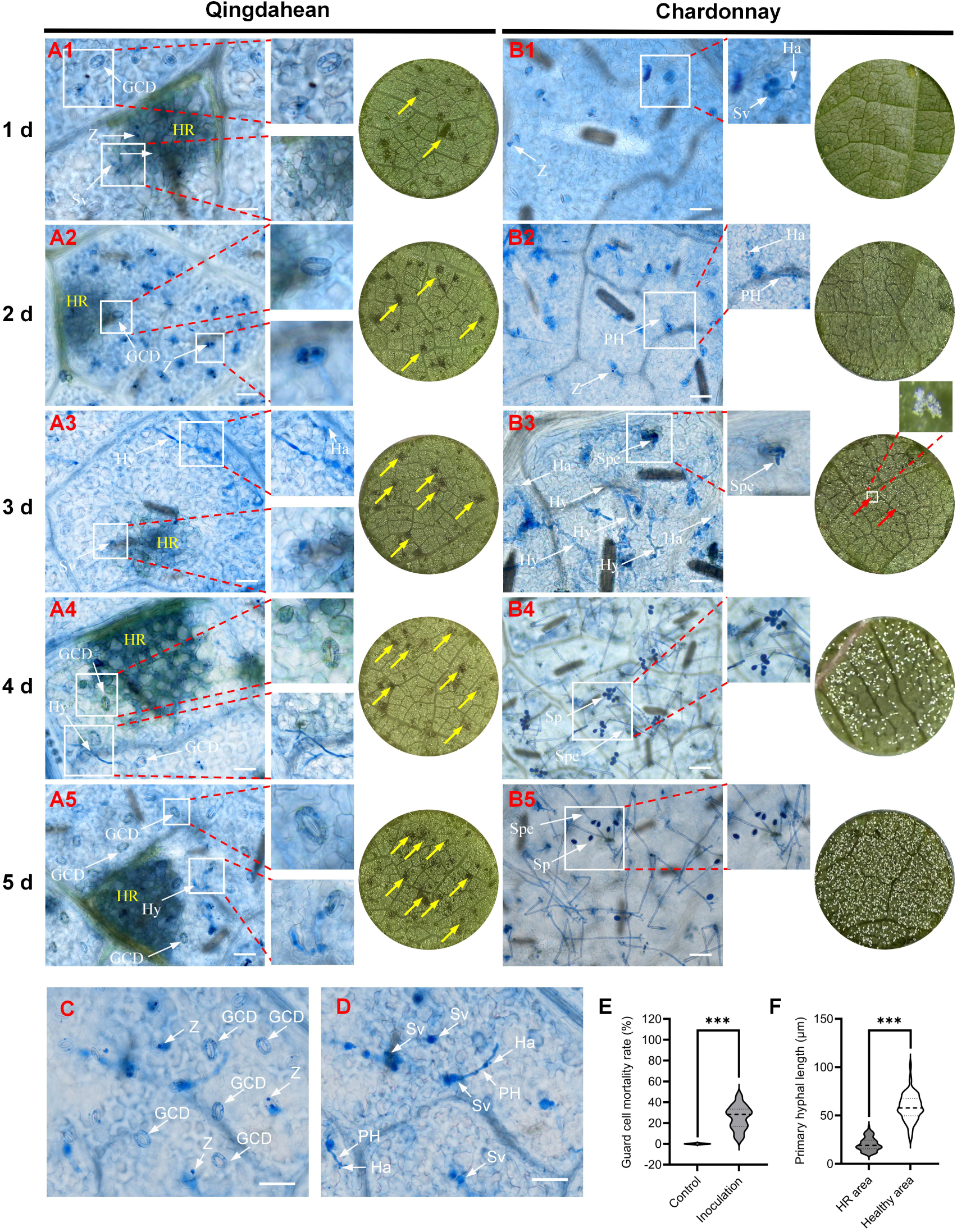
Host cell death induced by *P. viticola* in Qingdahean and Chardonnay leaves. **(A1-B5)** Representative images of *P. viticola* growth at different days post inoculation visualized by trypan blue staining. Representative photos of leaf discs are shown on the right of each panel. **(C)** Representative image of the surface layer depicting dead guard cells and the surrounding area in Qingdahean leaves at 2 DAI. **(D)** Representative image of the deeper layer shown in **C.** Blue color indicates stained *P. viticola* tissues and host cell death by trypan blue. Sp, sporangium; Spe, Sporangiophore; PH, Primary hypha; Hy, hypha; Z, encysted zoospores; Sv, substomatal vesicle; Ha, haustorium; GCD, guard cell death; HR, hypersensitive response; Yellow arrow, HR necrotic plaque; Red arrow, the sporangiophores of *P. viticola* emerging from the leaf discs. Inset, an enlarged view of the sporangiophores of *P. viticola*. Note not all the HR necrotic spots and sporangiophores of *P. viticola* in the leaf discs are indicated by arrows due to the large number. Leaf disk diameter, 8 mm, Scale bar, 50 μm. **(E-F)** Quantification of guard cell mortality rate **(E)** and primary hyphal length **(F)** in Qingdahean leaves at 2 DAI. Shown are violin plots and mean ± SE for n=28 regions from four independent leaf disks for **(E)** and n=115 primary hyphae lengths from four independent leaf disks for (F). The experiment was repeated three times with similar results (see Figure S13 for other independent sets of results). *\*\*\*P*<0.001, Student’s t-test.

Massive HR in Qingdahean has been observed from 1 DAI (Figure 4), so we performed a trypan blue staining assay to further investigate the effect of host cell death on the defense against *P. viticola*. It was shown that the HR necrosis spots and the death of some guard cells were already seen in Qingdahean leaves from 1 DAI (Figure 4A1) and increased over time (Figure 4). At 3 DAI, while Chardonnay leaves were extensively colonized by hyphae with a few sporangiophores protruding from the stomata, much fewer hyphae were seen inside of Qingdahean leaves (Figure 4A3, 4B3). At 4 and 5 DAI, Chardonnay leaves were largely covered by sporangium and sporangiophores (Figure 4B4, 4B5), but only a few sporangia were occasionally observed on Qingdahean leaves where there was no HR and guard cell death (Figure S7), indicating that the Qingdahean is not completely immune to *P. viticola*. Note, leaves that were not inoculated with *P. viticola* were used as a control and showed no symptom of cell death at all (Figure S8).

To learn more about how guard cell death and HR affected the *P. viticola* infection in Qingdahean, we conducted detailed quantification at 2 DAI, a time when hyphae did not spread widely and were easy to measure. A significant portion of guard cell death was observed in Qingdahean leaves at 2 DAI (Figure 4E), and the formation of substomatal vesicles and primary hyphae were observed mainly under healthy stomata (Figure 4C, 4D), suggesting that guard cell death is efficient in warding off *P. viticola* invasion. Similarly, in the HR necrotic area, the formation of substomatal vesicles was mainly found in the outskirt area (Figure S9). Quantification of the length of primary hyphae indicates that the HR response significantly suppressed the development of primary hyphae (Figure 4F). As a result, the development of primary hyphae and the formation of haustoria were both significantly suppressed in Qingdahean compared to Chardonnay (Figure S10). Taken together, these results suggest that cell death of guard cells and other host cells is a key defense strategy for *V. riparia* against *P. viticola*.

### Malonaldehyde positively regulates grapevine resistance against *P. viticola*

Since the resistance of Qingdahean against *P. viticola* occurred from the initial stage of stomatal targeting, we investigated the basal levels of H_2_O_2_, MDA, ABA, and SA to find clues about the resistance mechanism at the molecular level. The measurement results indicated that the MDA content in Qingdahean leaves was significantly greater than that in Chardonnay, but there were no differences observed in the contents of H_2_O_2_, ABA, and SA (Figure 5A, 5B, S11), which leads us to further investigate the role of MDA in grapevine resistance to DM.

**Figure 5.**
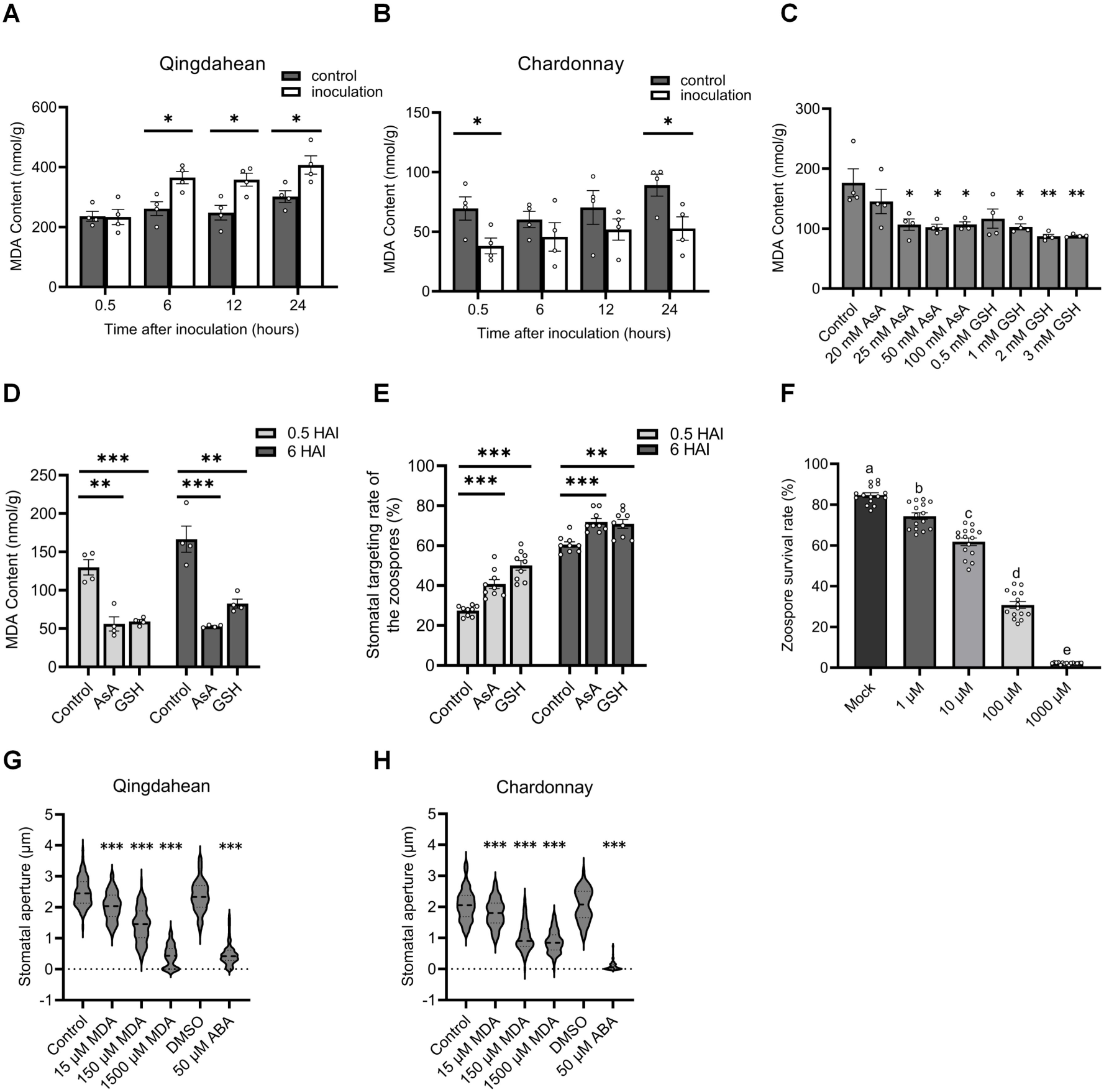
Effects of MDA on grapevine and *P. viticola*. **(A-B)** MDA content in Qingdahean and Chardonnay leaves at different HAI. **(C)** MDA content in Qingdahean leaf discs after 24 h treatment of AsA and GSH. **(D, E)** MDA content **(D)** and stomatal targeting rate of zoospores **(E)** at 0.5 and 6 HAI in Qingdahean leaf discs treated with 25 mM AsA and 2 mM GSH for 24 hours. Shown are data points and the mean ± SE for n=4 in **(A, B, C and D)** and for n=9 over four independent leaf disks in **(E)**. **(F)** Quantification of *P. viticola* zoospore viability shown in **(Figure S12)**. Zoospore survival rate was quantified as the ratio of the number of zoospores stained by FDA to the total number of zoospores. Shown are data points and means ±SE for n=15 images. Different letters indicate statistical significance (*P*<0.05, ANOVA with Duncan test). **(G, H)** Stomatal aperture size in Qingdahean and Chardonnay leaves after 2 h MDA treatment. Shown are individual data points and mean ± SE for n=100 stomata over four independent leaf disks. Control, pure water; 50 μM ABA treatment was used as a positive control to induce stomatal closure. The experiment was repeated three times with similar results (see Figure S13 for other independent sets of results). \**P*<0.05, \*\**P*<0.01, \*\*\**P*<0.001, Student’s t-test compared with Control.

Infection of *P. viticola* increased MDA concentration in Qingdahean but not in Chardonnay (Figure 5A, 5B). It is known that MDA is a principal product of lipid peroxidation of cell membranes and antioxidants including ascorbic acid (AsA) and glutathione (GSH) are effective in decreasing MDA content in plants (Nahar et al. 2015; Akram et al. 2017; Biswas and Mano 2021; Jalili et al. 2023; Kanwal et al. 2024). As shown in Figure 5C and 5D, treatment of AsA and GSH decreased the basal and *P. viticola* -induced MDA content in Qingdahaan, which significantly enhanced the stomatal targeting of *P. viticola* (Figure 5E). The effect of MDA on the viability of *P. viticola* zoospores was further investigated by fluorescein diacetate (FDA) staining assay. The results indicated that the viability of *P. viticola* was significantly decreased by MDA treatment in a dose-dependent manner (Figure 5F, S12). In addition, MDA itself triggered stomatal closure in a dose-dependent manner (Figure 5G, 5H). These results together suggest that the increased MDA content in Qingdahean leaves contributes to their resistance to downy mildew.

## DISCUSSIONS

In order to identify grapevine cultivars resistant to DM, the current study assessed 29 global cultivars from 7 species and observed that the European grapevine Chardonnay exhibited the highest number of sporangia per leaf disc, whereas the Muscadine grapes Fry, Granny Val, American hybrids 110R, Beta, SO4, 1103P, 5BB, and Qingdahean displayed the lowest number of sporangia (Figure S1B). These results are in line with previous reports showing that European grapevines are generally susceptible to downy mildew, while American and Muscadine grapes exhibit significant resistance to DM (Zhao et al. 2019; Heyman et al. 2021; Guo et al. 2024). The novel evaluated cultivars are important germplasms for grapevine improvement. Chardonnay and Qingdahean were selected as susceptible and resistant cultivars to further understand the resistance mechanism against *P. viticola*. A major defense phenotype observed in Qingdahean is the occurrence of many necrotic patches on leaves upon inoculation, a hallmark of HR. Microscopy analysis further revealed intensive stomatal defenses against *P. viticola* that have been overlooked so far.

*Plasmopara viticola* is specialized to internalize into leaves and disperse spores through stomata. Previous studies have observed stomatal opening in the dark and deregulation induced by *P. viticola* infection after 3 DAI in a susceptible *V. vinifera* cultivar Marselan, which is proposed to facilitate the emergence of sporangiophores and pathogen dissemination (Allègre et al. 2007). However, the early stomatal defenses within hours, such as stomatal closure, inhibition on the stomatal targeting and penetration, and the different response between stomata with and without pathogen remains largely unknown. By sophisticated histochemical studies of the susceptible grapevine Chardonnay and resistant grapevine Qingdahean, the current results provide evidence that stomatal defense against *P. viticola* contains at least 4 layers (Figure 6): 1) interference in stomatal targeting seen in the Qingdahean; 2) stomatal closure seen in both cultivars; 3) interference in stomatal invasion seen in Qingdahean; 4) guard cell death seen in Qingdahean. Regarding the molecular mechanism, previous studies from bacteria and fungi have shown that molecules both from microbes and the host such as bacterial flagellin, fungal cell wall components chitin and chitosan, phytocytokine Peptide elicitor 1, function as triggers inducing stomatal closure and guard cell death (Melotto et al. 2006; Zheng et al. 2018; Ye et al. 2020). In terms of *P. viticola*-induced stomatal closure, cell wall components may be the candidate molecules, as mixtures of β-1→3-linked glucans, major components of oomycete, have been shown to induce stomatal closure in grapevine (Allègre et al. 2009). So far, there are over 30 loci associated with *P. viticola* resistance (*Rpv*), and *Rpv5* and *Rpv6* are identified from *V. riparia* cultivar Gloire de Montpellier (Marguerit et al. 2009; Koledenkova et al. 2022; Ricciardi et al. 2024). However, only *Rpv1* and *Rvp3* are characterized to gene level with functional validation in the past research showing that several TIR-NB-LRR (TNL) genes are responsible for the host cell death and resistance against *P. viticola* (Feechan et al. 2013; Foria et al. 2020). Future work is necessary to test whether these TNL genes play a role in the host cell death observed here including guard cell death.

**Figure 6.**
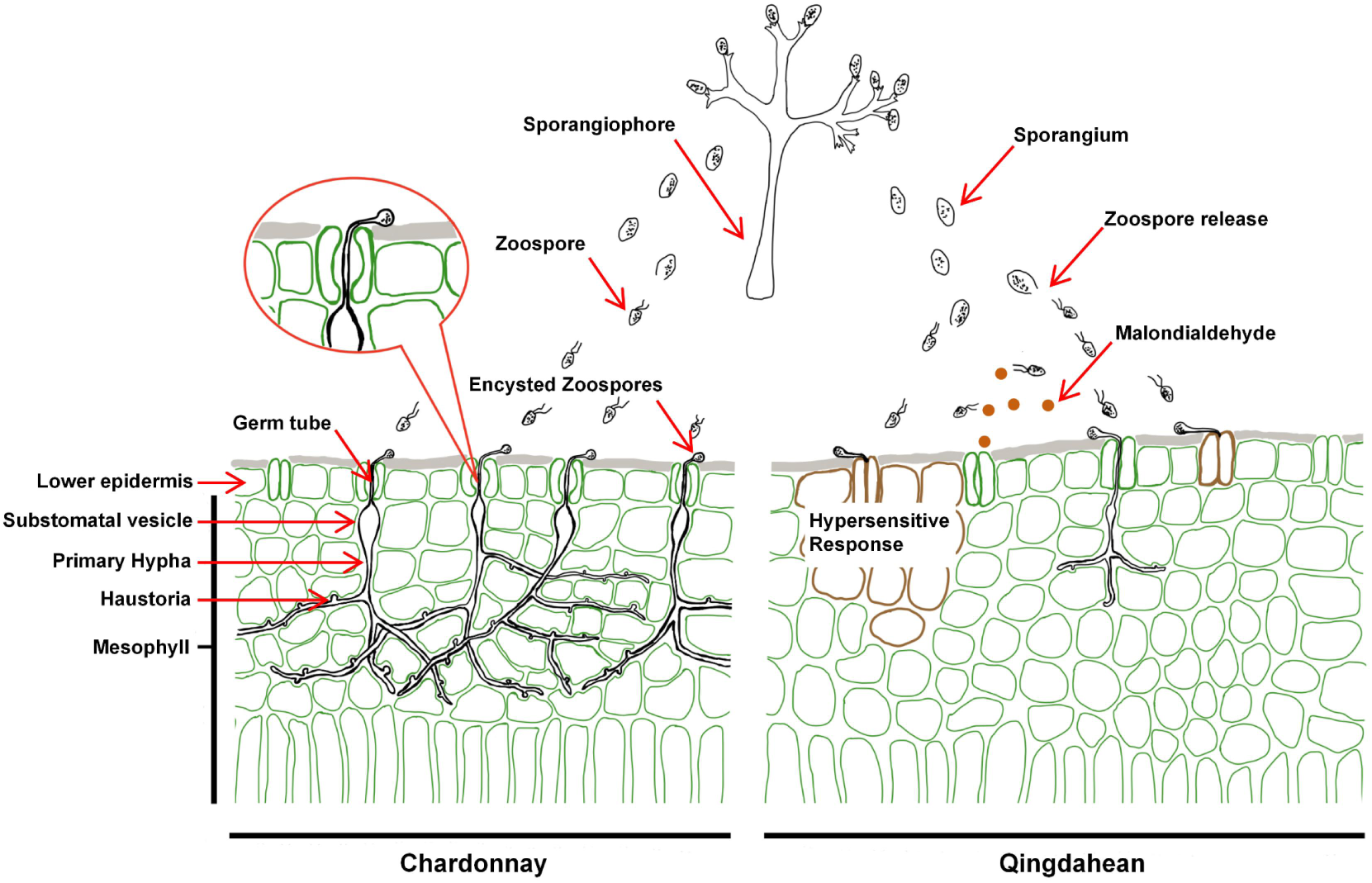
Proposed layered stomatal immunity against *P. viticola* infection in grapevine leaves. By comparison of *P. viticola* infection in Chardonnay and Qingdahean, the following defense responses at the stomatal level was proposed: 1) interference in stomatal targeting of zoospores in the Qingdahean; 2) stomatal closure seen in both cultivars; 3) interference in stomatal invasion of germ tubes in Qingdahean; 4) guard cell death in Qingdahean. MDA molecules released from stomata repel the zoospores, interfering the stomatal targeting process in the Qingdahean.

Malondialdehyde is a highly reactive and diffusible product of polyunsaturated fatty acid oxidation, predominantly occurring in cellular membranes, in plant cells involved in biotic and abiotic stress responses (Mohamed et al. 2012; Morales and Munné-Bosch 2019; Biswas and Mano 2021). According to reports, MDA has a concentration ranging from nmol g⁻¹ fresh weight (FW) to μmol g⁻¹ FW in grapevine (Mozafari et al. 2018; Li and Wang 2021; Jiao et al. 2023; Yang et al. 2023), and is able to strengthen the resistance of grapevines to DM by activating several signaling pathways and defense mechanisms (Heyman et al. 2021). Our results showed that MDA content was significantly higher in Qingdahean than that in Chardonnay at resting status and was further increased in Qingdahean following inoculation with *P. viticola*, that decreased MDA content by antioxidants facilitated the stomatal targeting of *P. viticola*, and that MDA had an inhibitory effect on zoospore viability (Figure 5). These results lead us to hypothesize that MDA, leaking from stomatal openings, functions as a toxic compound to repel zoospores resulting in the impaired stomatal targeting rate seen in Qingdahean. On the other hand, *P. viticola* may have strategies to tolerate the toxicity of MDA in grapevine, as seen that stomatal targeting rate was still substantial in Qingdahean. Reactive products of lipid peroxidation of cell membranes are known signals required for stomatal closure under different stresses (Islam et al. 2016; Islam et al. 2020; Biswas and Mano 2021). Since MDA induced stomatal closure in grapevines (Figure 5G), it is possible that MDA is involved in the rapid stomatal responses to *P. viticola* in Qingdahean. H_2_O_2_, ABA, and SA are important signaling molecules involved in stomatal defenses and other immune responses (Kim Khiook et al. 2013; Yu et al. 2016; Bharath et al. 2021). Though our results did not find differences in the basal content of H_2_O_2_, ABA, and SA in both cultivars, it remains to be clarified whether their recruitment induced by *P. viticola* contributes to the resistance differences of the two cultivars.

On the pathogen side, it seems that *P. viticola* has evolved strategies to overcome stomatal immunity, as seen in the successful infection of Chardonnay. Interestingly, the stomata not targeted by the pathogens close rapidly while the targeted ones did not in Chardonnay (Figure 2), suggesting that there are signals from *P. viticola* manipulating stomatal movement. In guard cell-bacterium interaction, it is well-known that bacteria secrete toxins and effectors to manipulate stomatal aperture and host cell fate eventually breaking stomatal immunity (Melotto et al. 2006; Hou et al. 2021). RXLR (Arg-any amino acid-Leu-Arg) and CRNs (crinkling and necrosis induced protein) effectors are two classes of major effectors secreted by pathogenic oomycetes, including *P. viticola* to overcome host immunity (Ma et al. 2020; Liu et al. 2021; Xiang et al. 2021). On the other hand, it is known that effectors are mainly secreted after haustoria formation during oomycete infection, which is a post-stomatal invasion event (Boevink et al. 2020; Bozkurt and Kamoun 2020). The current study raises interesting questions whether these effectors are secreted during stomatal invasion and involved in the suppression of stomatal immunity.

In summary, the present study provides details of layered stomatal defenses of grapevine previously ignored against *P. viticola*, paving the way for future work on stomatal immunity against pathogenic oomycete in general.

## MATERIALS AND METHODS

### Grapevines and pathogen

The *Vitis* germplasms consist of 2 genera, 7 species, and 29 cultivars originating from various areas across the globe (Table S1) and were grown in a glasshouse at a temperature of 28–20°C (day:night). Standard management procedures, such as cultivation, watering and fertilization were applied to all plants.

For mechanism study of Chardonnay and Qingdahean resistance to downy mildew, the grapevines were grown in pots (60% peat moss, 20% vermiculite, 20% perlite) in a growth chamber with light intensity of 200 μmol·m^-2^·s^-1^, temperature of 28±2°C, humidity 85±5%, and a 16 h light/8 h dark regime.

*Plasmopara viticola* was maintained on detached Chardonnay leaves in a petri dish with bottom covered by fully wetted filter paper incubated in a growth chamber with light intensity of 55 μmol·m^-2^·s^-1^, temperature of 19 ± 2°C, humidity 85 ± 5%, and a 12-hour light/dark regime.

### Pathogen inoculation

For disease evaluation of the 29 grapevine cultivars, the inoculation was performed according to previous study (Paineau et al. 2022; Guo et al. 2024). Each leaf disc (8 mm) was inoculated with 20 μL *P. viticola* suspension in water of 1 × 10^5^ sporangia/mL on the abaxial surface. Excess suspension was removed after 24 h incubation. At 10 DAI, the symptom of each leaf discs was imaged and the number of sporangia was counted.

For the resistance mechanism study of Qingdahean and Chardonnay, the leaf discs (6 mm) were inoculated by floating on 200 μL *P. viticola* suspension of 1 × 10^5^ sporangia/mL with the abaxial surface facing the suspension. The leaf discs were moved away from the suspension after 24 h and incubated in a petri dish having a fully wet filter paper inside with abaxial surface facing up to develop symptom.

### Blankophor and Trypan blue staining

Each leaf disc (6 mm) was floated on 200 μL *P. viticola* suspension of 1 × 10^5^ sporangia/mL with the abaxial surface facing the suspension. The leaf discs were taken out at indicated time point for staining. For blankophor staining, each leaf disc was surface-stained with the blankophor solution (A 1‰ (v/v) Blankophor stock solution was prepared in water and the work solution was a 5% dilution of the stock in water or in 15%(w/v) KOH), as described previously (Díez-Navajas et al. 2007), subsequently mounted onto a glass slide, and allowed to equilibrate for a period of 2 min at ambient temperature conditions. Thereafter, the samples were immediately visualized under a fluorescence microscope (Eclipse Ti2-E, DS- 10, Nikon Corporation, Japan) with a DAPI filter, which has an excitation wavelength of 360-390 nm and an emission range of 430-490 nm. The stomatal targeting rate was calculated as the ratio of the number of stomata having encysted spores to the number of all stomata in the entire field of view.

For trypan blue staining, the leaf discs were submerged in trypan blue solution (0.67 mg/ml trypan blue in 1:1:1:1:6 [vol:vol] solution of water:glycerol:lactic acid:phenol:ethanol, respectively) and boiled for 15 min (Gao et al. 2016). After overnight cooling, the leaf discs were decolorized in chloral hydrate (2.5 g/mL). Once the leaf discs became transparent, they were imaged in a microscope (Eclipse Ci-L, DS- Fi3, Nikon Corporation, Japan). Stomatal invasion rate was calculated as the ratio of the number of stomata having substomatal vesicles to the number of stomata targeted by zoospores. The length of the primary hyphae was measured by ImageJ (ImageJ, 1.52V, Wayne Rasband, NIH, USA). Guard cell death rate was calculated as the ratio of number of stained guard cells to the number of all stomata in the entire field of view.

### Stomatal aperture measurement

Leaf discs (6 mm) were floated on water with their adaxial surface upward in the light (600 µmol·m⁻²·s⁻¹) for 2 h to open stomata (Ye et al. 2020). Then the water was replaced with 200 μL *P. viticola* suspension of 1 × 10^5^ sporangia/mL and incubated for the indicated time followed by imaging using a microscope (Eclipse Ci-L, DS-Fi3, Nikon Corporation, Japan). The aperture was measured with Image J (ImageJ, 1.52V, Wayne Rasband, NIH, USA).

### Quantification of H_2_O_2_, MDA, ABA, and SA content

Fully-expended grapevine leaves were used to quantify MDA, H_2_O_2_, ABA, and SA. Leaf samples were collected and rapidly frozen in liquid nitrogen. Subsequently, they were ground into a fine powder using a mortar. The resulting powder was then transferred into 25 mL centrifuge tubes and stored in a - 80°C freezer before use. The MDA and H_2_O_2_ content were quantified using a Malondialdehyde Content Detection Kit and Hydrogen Peroxide Content Detection kit (Shanghai Acmec Biochemical Co., Ltd. China) according to the manufacturer instruction.

The hormones ABA and SA were quantified according to previous methods (Yao et al. 2022). All the detections were performed on a Vanquish UHPLC system combined with a TSQ Altis MS/MS system (Thermo Scientific, USA). ACQUITY UPLC@HSS T3 column (150 mm × 2.1 mm, 1.8 μm particle size, Waters) was used for the separation of samples with the column oven at 35°C. The mobile phase was composed of 0.1% formic acid in water (A) and 0.1% formic acid in ACN (B) with the flow rate at 0.2 mL/min. A linear gradient elution program with the following proportions (v/v) of solvent B was applied: 0-1 min at 10%, 1-13 min from 10% to 60%, 13-14 min from 60% to 95%, 14-17 min at 95%, 17-18.1 min from 95% to 10%, 18.1-20 min at 10%, giving a total run time of 20 min. The injection volume was 2 µL. Salicylic acid and ABA were monitored by multiple reaction monitoring (MRM) in a positive/negative mode of Heated-electrospray ionization(H-ESI). The ion source conditions were as follows: positive ion capillary, 4000 V; negative ion capillary, 3000 V; sheath gas, 35 arb; aux gas, 10 arb; ion transfer tube temperature, 350°C; vaporizer temperature, 350°C. After optimizing the MRM parameters, the precursor ion m/z of SA under negative mode was 136.8, while the main specific products ion m/z were 64.883 and 92.883 with the collision energy of 28.71 and 16.02 V, respectively. The precursor ion m/z of ABA under negative mode was 262.967, while the main specific products ion m/z were 152.967 and 219.05 with the collision energy of 10.41 and 14.45 V, respectively. The RF lens was set at 42V for SA and 44 V for ABA.

### Zoospore viability assay

A suspension of *P. viticola* of 1 × 10^5^ sporangia/mL was incubated for 2 h to fully release the zoospores. The suspension was treated with MDA solutions at indicated concentrations in the dark at 19°C for 30 minutes, followed by staining with FDA (2 µg/mL) for 2 minutes. The zoospores stained by FDA were imaged under a fluorescence microscope (Eclipse Ti2-E, DS-10, Nikon Corporation, Japan) having a FITC filter with an excitation wavelength of 465-495 nm and an emission range of 512-558 nm. The viability of the zoospores was calculated as the ratio of the number of zoospores stained with FDA to the total number of zoospores in the field of view.

### Determination of chlorophyll content and photosynthetic parameters

Chardonnay and Qingdahean grown in pots were entirely spray-inoculated with *P. viticola* suspension of 1 × 10^5^ sporangia/mL. Leaves from the 4th to 7th leaves (counted downwards from the top) were used for all measurement at 1, 3, 5, 7, and 10 DAI according to previous methods (Nogueira Júnior et al. 2019). The chlorophyll content was quantified using a SPAD-502 plus measuring device (Konica Minolta Sensing Inc., Japan), with random sections being chosen on each leaf. Fv/Fm, ETR, and PhiPS2 were measured by the LI-600 fluorescence-stomatal measurement apparatus (LI-COR in Lincoln, NE, USA).

Net photosynthetic rate (Pn), stomatal conductance (Gsw), intercellular CO_2_ concentration (Ci), and transpiration rate (E) were quantified by the LI-6800 portable photosynthesis equipment (LI-COR, Lincoln, NE, USA) according to the manufacturer instruction. The flow rate, leaf temperature, relative humidity, light intensity and CO_2_ concentration were kept constant at 500 μmol s^−1^, 26 °C, 60% (Pa/Pa), 1500 μmol m^-2^ s^-1^ and 400.0 µmol mol⁻¹, respectively.

## ACKNOWLEDGEMENTS

We thank PKU-IAAS Mass Spectrometry Platform for their technical support in quantification of SA and ABA. W.Y. laboratory is supported by Taishan Scholars Program of Shandong Province and Shandong Provincial Science and Technology Innovation Fund.

## AUTHOR CONTRIBUTIONS

W.Y. designed the experiments. W.Z., W.J. J.M., X.L., G.Q. performed all experiments and analyzed all data. W.Z., M.S.R, N.A. and W.Y. interpretated results. W.Z., M.S.R. and W.Y. wrote the manuscript.

## Supplemental Table and Figures

Table S1. Information of the 29 Cultivars

Figure S1. Global distribution of 29 grape cultivars and their resistance to downy mildew

Figure S2. Leaf morphology of different grape cultivars used in this study

Figure S3. Symptoms of leaf discs of different grape cultivars at 10 days post inoculation

Figure S4. Representative images of symptoms of Chardonnay and Qingdahean whole plants at different days post inoculation

Figure S5. Changes in chlorophyll content and photosynthetic parameters of Qingdahean and Chardonnay leaves at different days post inoculation

Figure S6. The stomatal density of Qingdahean and Chardonnay

Figure S7. Rare sporulation event on Qingdahean leaves infected by *P. viticola* at 4 DAI, as visualized by trypan blue staining

Figure S8. Images and trypan blue staining of Qingdahean and Chardonnay leaf discs at different days post-inoculation with pure water as the control treatment

Figure S9. Tissue showing hypersensitive response and adjacent healthy tissue in Qingdahean leaves at 2 DAI visualized by trypan blue staining

Figure S10. The development of primary hyphae and haustoria in Qingdahean and Chardonnay leaves at 2 DAI

Figure S11. Quantification of MDA, H_2_O_2_, ABA, and SA basal content of Qingdahean and Chardonnay leaves

Figure S12. FDA staining of *P. viticola* zoospores treated with MDA

Figure S13. The other two independent sets of results for Figure 1-Figure 5

