## Supplementary material for "Layered stomatal immunity contributes to resistance of *Vitis riparia* against downy mildew *Plasmopara viticola*": Table S1 and Figure S1-S13

**Table S1. Information of the 29 Cultivars**

| Serial No | Cultivars | Species | Country of Origin | Use |
|---|---|---|---|---|
| V01 | Fry | *V. rotundifolia* Michx. | USA | Table grape |
| V02 | Granny Val | *V. rotundifolia* Michx. | USA | Table grape |
| V03 | 110 R | *V. berlandieri* × *V. rupetris* | France | Root stock |
| V04 | Beta | *V. labrusca* L.× *V. riparia* | USA | Root stock |
| V05 | SO4 | *V. berlandieri* × *V. riparia* | Germany | Root stock |
| V06 | 1103P | *V. berlandieri* × *V. rupetris* | Italy | Root stock |
| V07 | 5BB | *V. berlandieri* × *V. riparia* | Austria | Root stock |
| V08 | Qingdahean | *V. riparia* | USA | Root stock |
| V09 | Yongyou No.1 | *V. vinifera* L. × *V. labrusca* L. | China | Table grape |
| V10 | Molixiang | *V. vinifera* L. × *V. labrusca* L. | China | Table grape |
| V11 | Summer black | *V. Labrusca* L. × *V. vinifera* L. | Japan | Table grape |
| V12 | Sauvignon Blanc | *V. vinifera* L. | France | Wine grape |
| V13 | Riesling | *V. vinifera* L. | Germany | Wine grape |
| V14 | Moldova | *V. Labrusca* L. × *V. vinifera* L. | Moldova | Table grape |
| V15 | Cabernet Sauvignon | *V. vinifera* L. | France | Wine grape |
| V16 | Nantaihutezao | *V. vinifera* L. × *V. labrusca* L. | China | Table grape |
| V17 | Queen Nina | *V. vinifera* L. × *V. labrusca* L. | Japan | Table grape |
| V18 | Beibinghong | *V. amurensis* × *V. vinifera* L. | China | Wine grape |
| V19 | Kyoho | *V. Labrusca* L. × *V. vinifera* L. | Japan | Table grape |
| V20 | Wagamichi | *V. vinifera* L. × *V. labrusca* L. | USA | Table grape |
| V21 | Red Globe | *V. vinifera* L. | USA | Table grape |
| V22 | Thompson Seedless | *V. vinifera* L. | Turkey | Table grape |
| V23 | Victoria | *V. vinifera* L. | Romania | Table grape |
| V24 | Shine Muscat | *V. Labrusca* L. × *V. vinifera* L. | Japan | Table grape |
| V25 | Muscat Hamburg | *V. vinifera* L. | U. Kingdom | Table grape |
| V26 | Gewürztraminer | *V. vinifera* L. | Italy | Wine grape |
| V27 | Manicure Finger | *V. vinifera* L. | Japan | Table grape |
| V28 | Cabernet Franc | *V. vinifera* L. | France | Wine grape |
| V29 | Chardonnay | *V. vinifera* L. | France | Wine grape |

**Figure S1. Global distribution of 29 grape cultivars and their resistance to downy mildew**

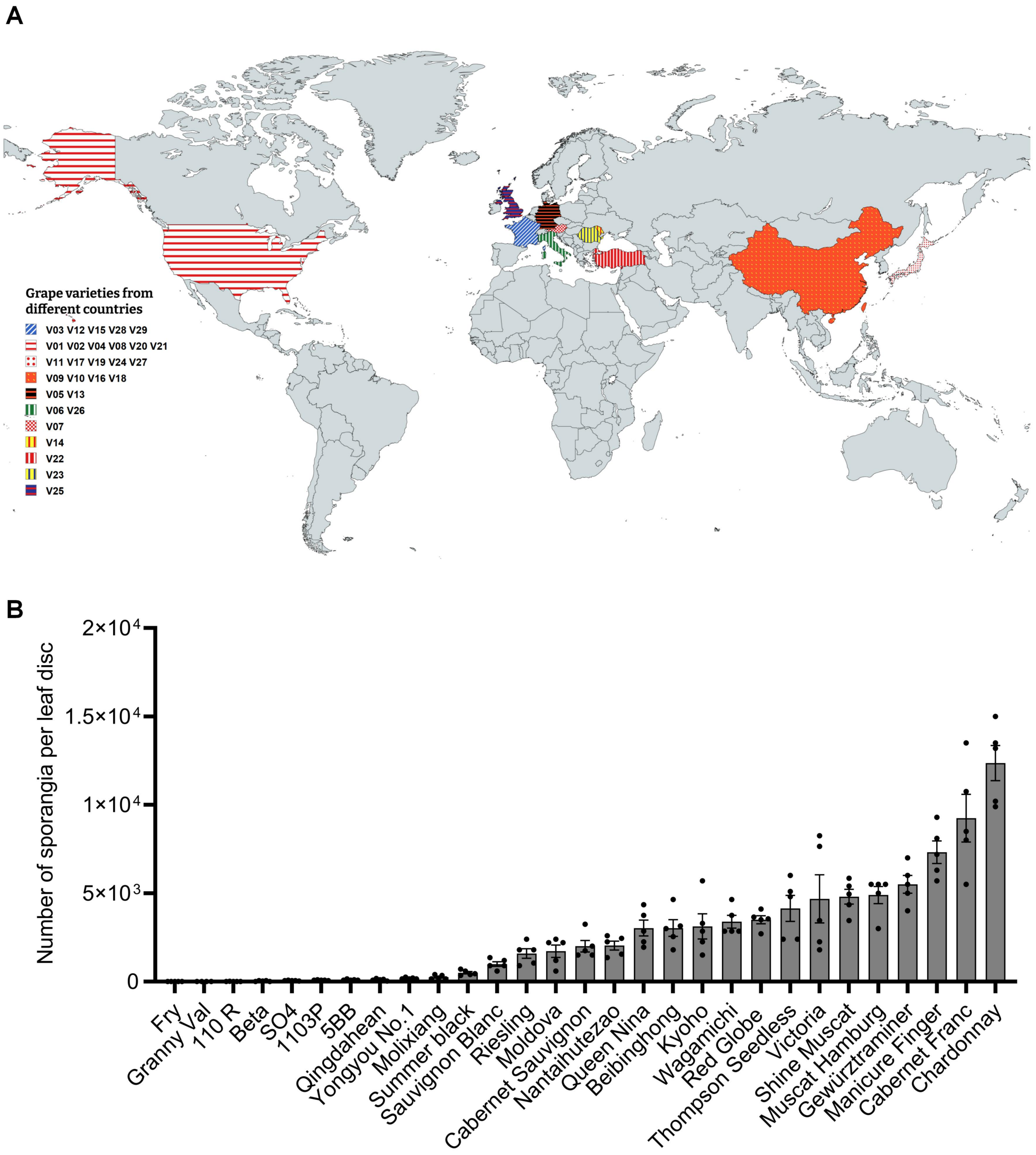


**(A)** Geographic distribution of the grapevines used in this study. Different color highlights the country position of grapevine cultivars. The map is generated by mapchart.net online. **(B)** The number of sporangia on the leaf discs of different grape cultivars. Each dot plot in the bar graph represents a replication. Shown are individual data points and mean ± SE for n=5 leaf disks over multiple independent experiments.

**Figure S2. Leaf morphology of different grape cultivars used in this study**

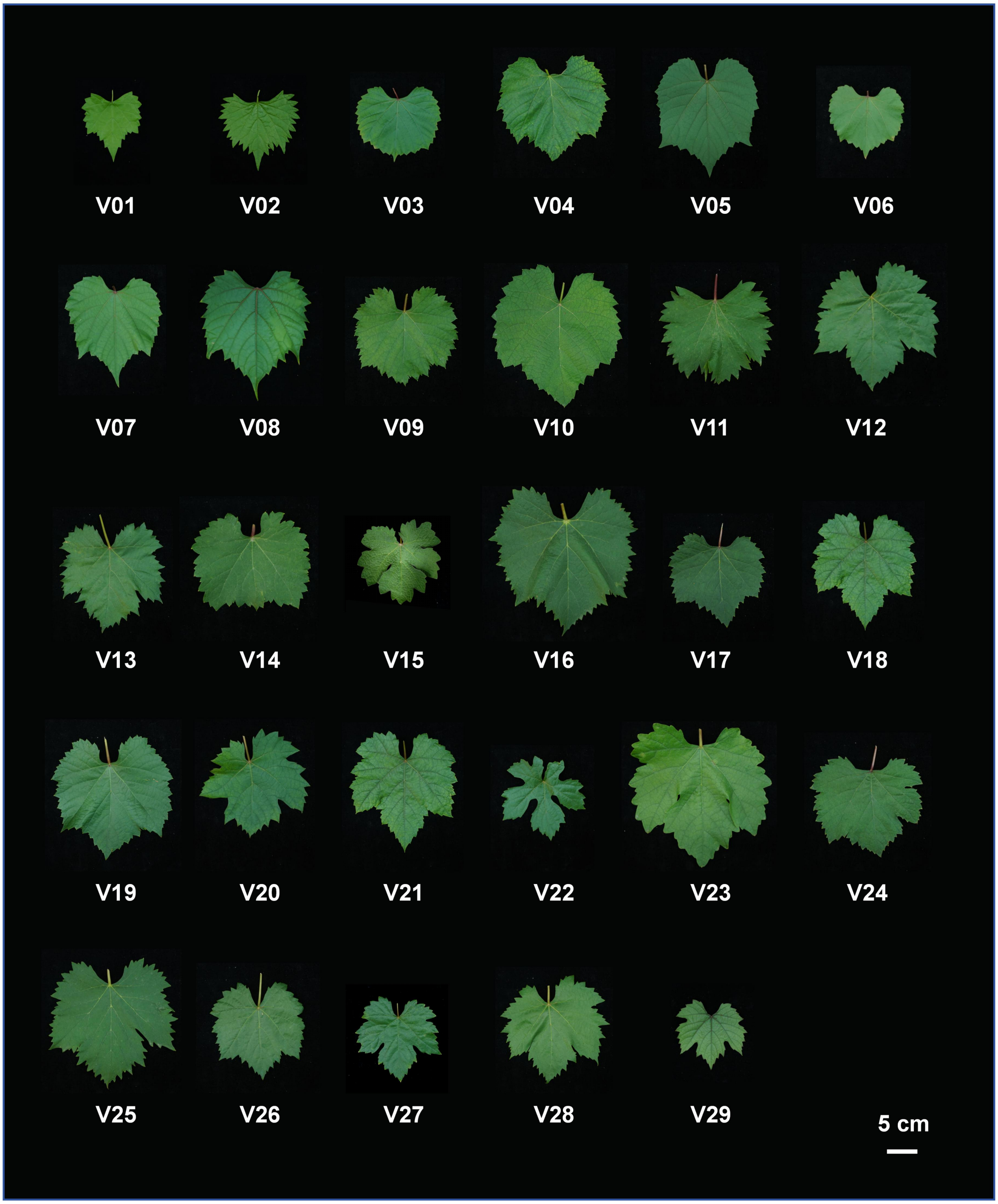

**Figure S3. Symptoms of leaf discs of different grape cultivars at 10 days post inoculation**

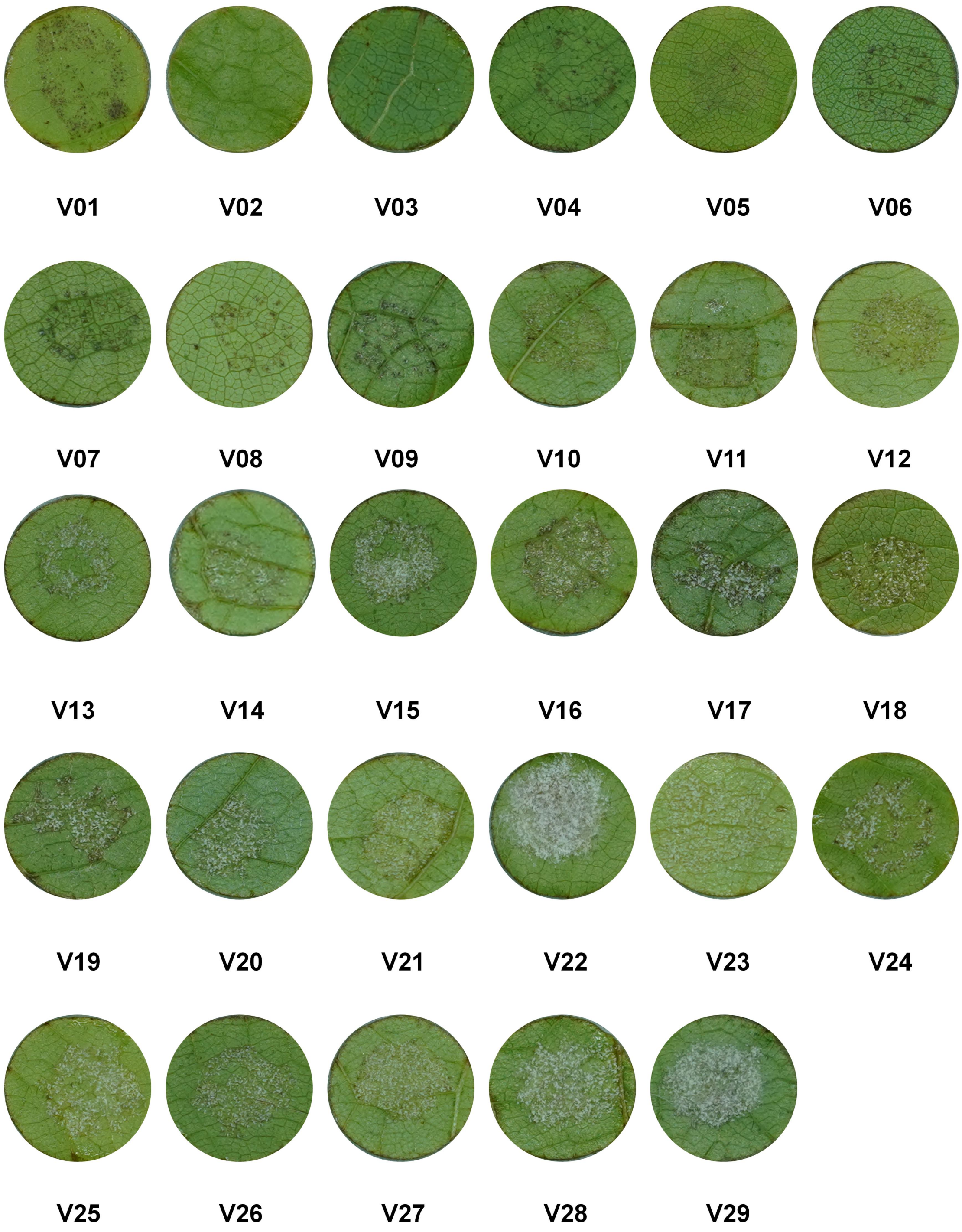

**Figure S4. Representative images of symptoms of Chardonnay and Qingdahean whole plants at different days post inoculation**

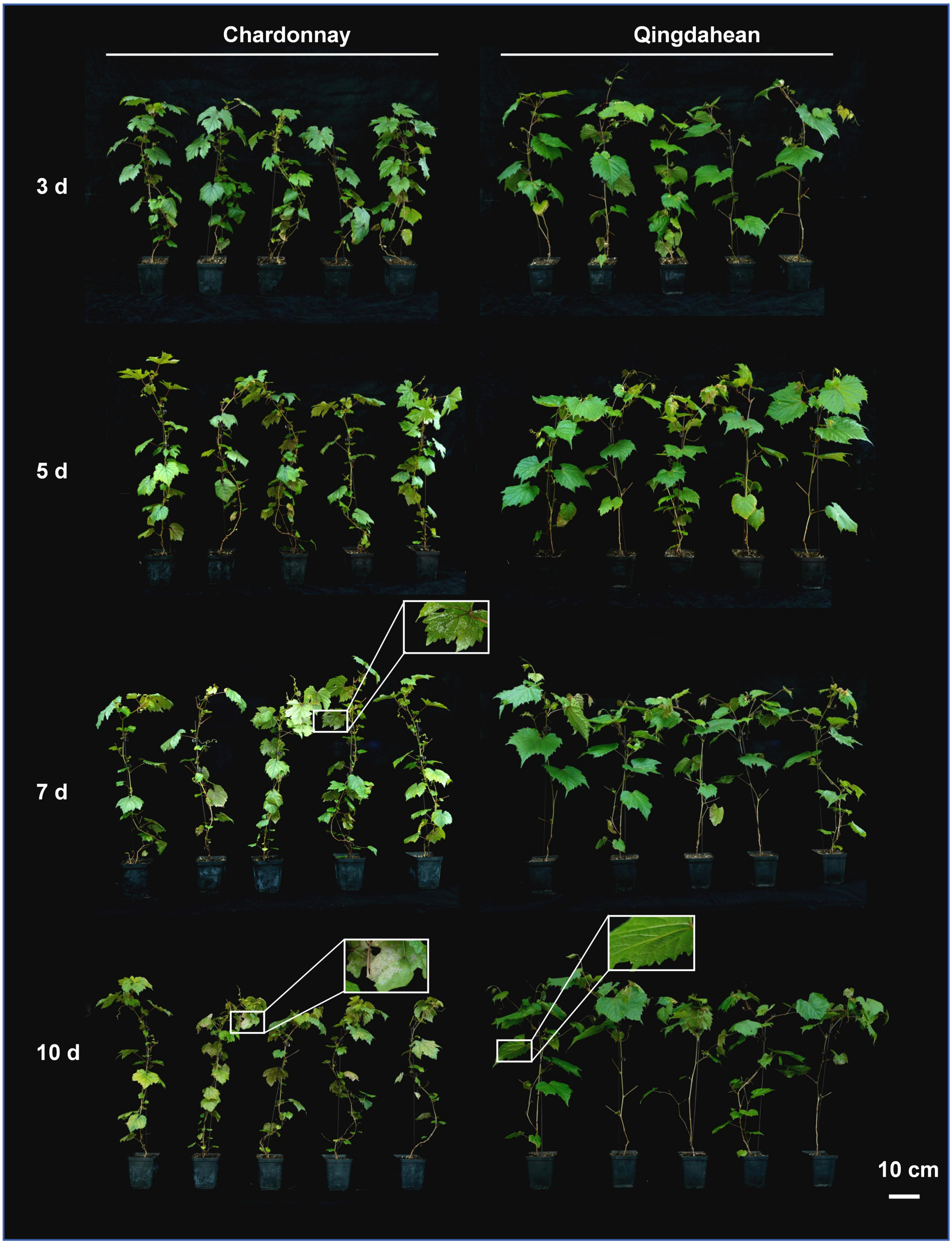


Inset, an enlarged view showing infection symptoms of *P. viticola* on the abaxial leaf surface.

**Figure S5. Changes in chlorophyll content and photosynthetic parameters of Qingdahean and Chardonnay leaves at different days post inoculation**

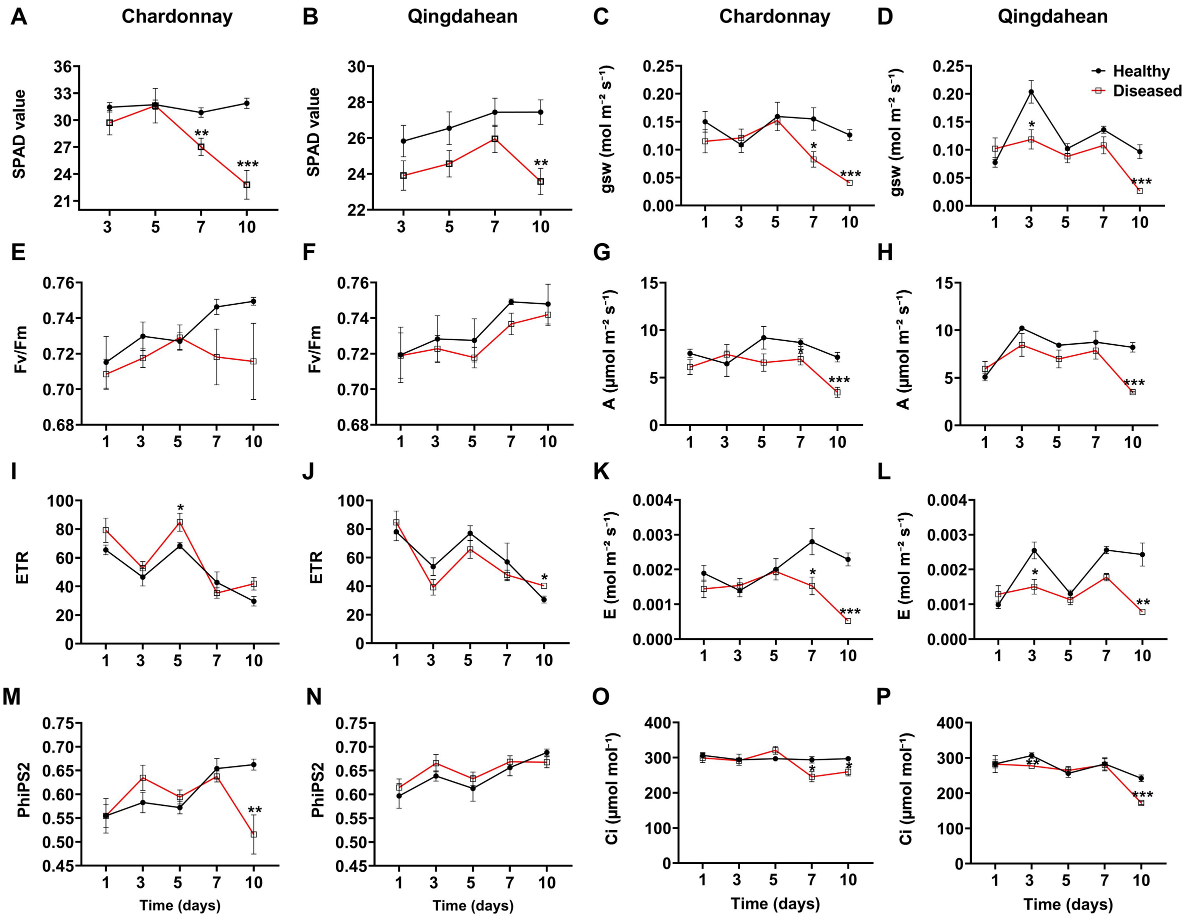


SPAD value, Soil Plant Analysis Development value; Fv/Fm, maximum photochemical efficiency; ETR, Electron transport rate; PhiPS2, actual photochemical efficiency; gsw, stomatal conductance; A, Net photosynthetic rate; E, Transpiration rate; Ci, intercellular carbon dioxide concentration. Shown are individual data points and mean ± SE obtained from replicates of five plants. **P*<0.05, ***P*<0.01, ****P*<0.001, Student's t-test.

**Figure S6. The stomatal density of Qingdahean and Chardonnay**

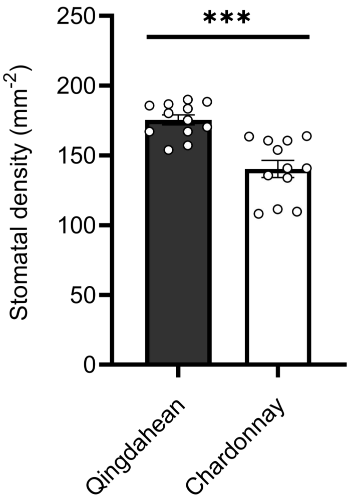


The figures show the mean ± SE of stomatal density in 12 randomly selected areas from four leaves of Qingdahean and Chardonnay, respectively. ****P*<0.001, Student's t-test.

**Figure S7. Rare sporulation event on Qingdahean leaves infected by *P. viticola* at 4 DAI, as visualized by trypan blue staining**

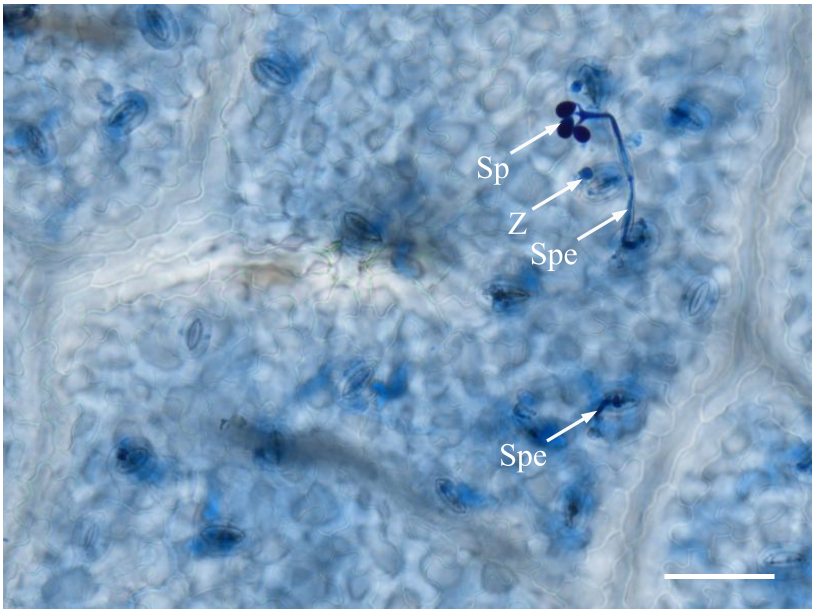


Scale bar, 50 μm; Sp, sporangium; Spe, Sporangiophore; *Z*, encysted zoospores.

**Figure S8. Images and trypan blue staining of Qingdahean and Chardonnay leaf discs at different days post-inoculation with pure water as the control treatment**

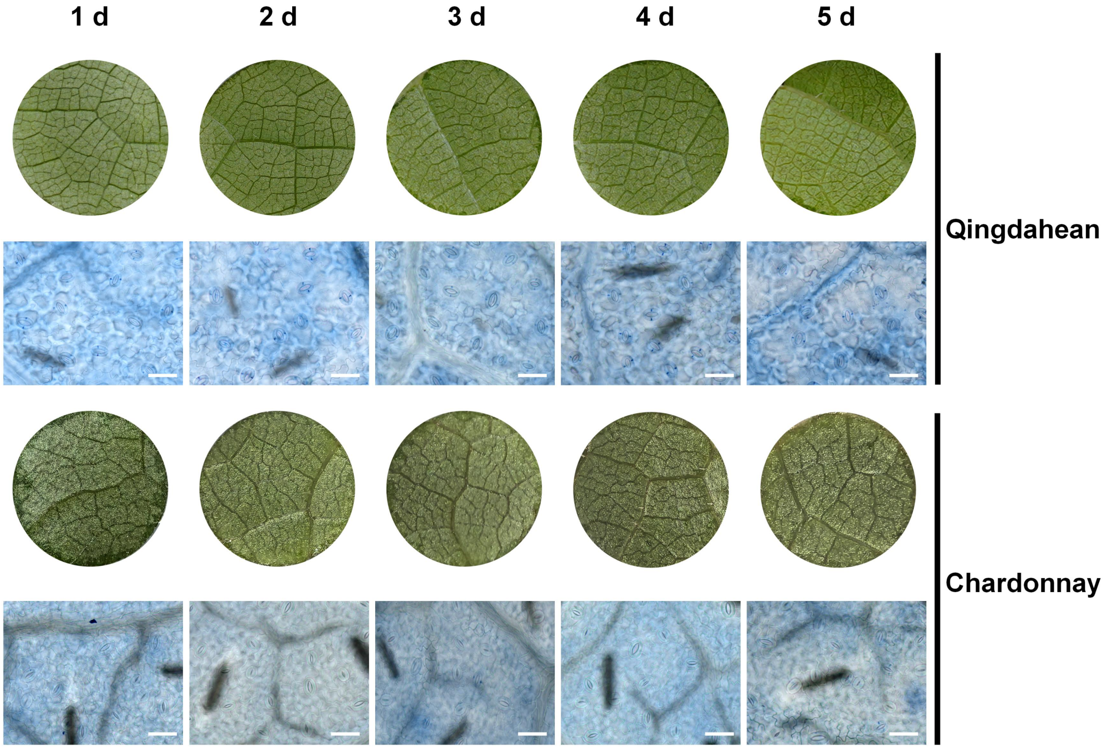


Leaf disk diameter, 8 mm; Scale bar, 50 μm.

**Figure S9. Tissue showing hypersensitive response and adjacent healthy tissue in Qingdahean leaves at 2 DAI visualized by trypan blue staining**

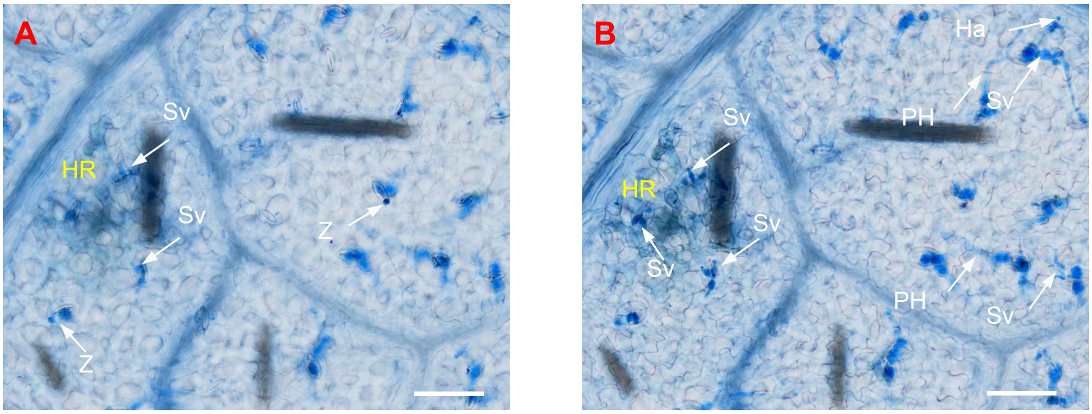


**(A)** Representative image of the hypersensitive response (HR) area and the surrounding healthy mesophyll cell area at the surface level in Qingdahean leaves at 2 DAI. **(B)** Representative image of the deeper layer shown in A.

Scale bar, 50 µm. Ha, haustorium; HR, hypersensitive response; PH, Primary hypha; Sv, substomatal vesicle; Z, encysted zoospores.

**Figure S10. The development of primary hyphae and haustoria in Qingdahean and Chardonnay leaves at 2 DAI**

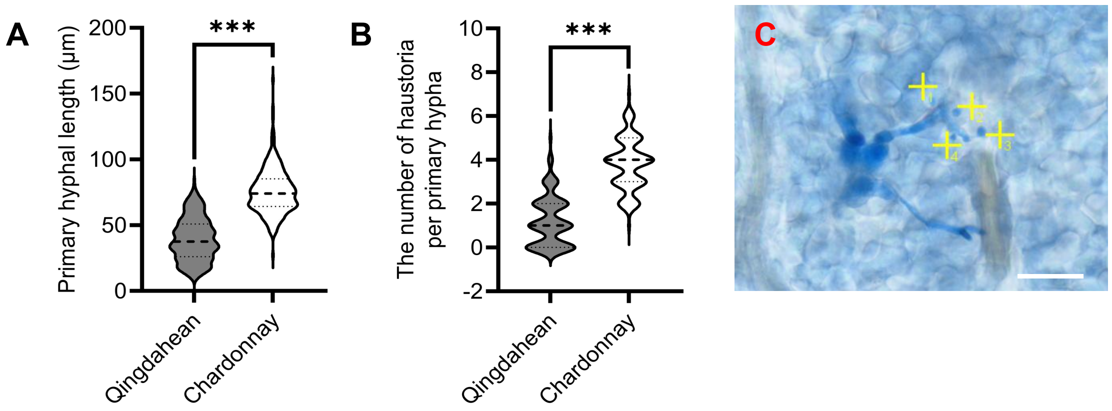


**(A)** Quantification of the length of primary hypha. Shown are the violin plots and mean ± SE for n=260 regions sampled from four independent leaf disks. ****P*<0.001, Student's t-test. **(B)** Quantification of the number of haustoria per primary hyphae. Presents the violin plots and mean ± SE for n > 100 primary hyphae lengths sampled from four independent leaf disks. ****P*<0.001, Student's t-test. **(C)** Representative images showing the quantification of the number of haustoria per primary hypha; Scale bar, 20 μm.

**Figure S11. Quantification of MDA, $H_2O_2$, ABA, and SA basal content of Qingdahean and Chardonnay leaves**

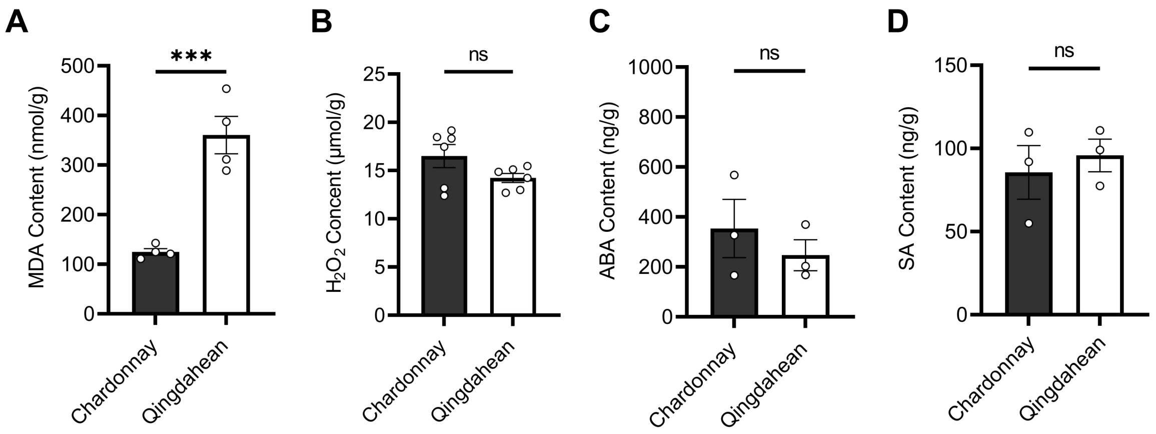


Quantification of MDA **(A)**, $H_2O_2$ **(B)**, ABA **(C)**, and SA **(D)** basal content of Qingdahean and Chardonnay leaves. Shown are individual data points and mean ± SE for n=4 plants for **(A)**, n=6 plants for **(B)**, and n=3 plants for **(C, D)**. The experiment was repeated three times with similar results. ***$P$<0.001, Student's t-test. ns, not significant.

**Figure S12. FDA staining of *P. viticola* zoospores treated with MDA**

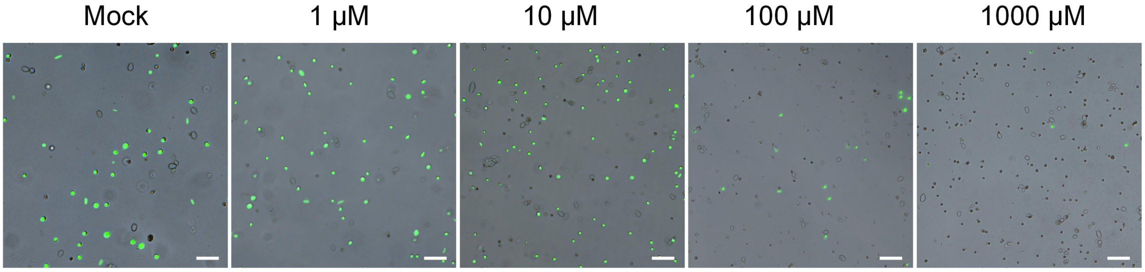


Representative overlay images of the bright and fluorescence view were shown. Scale bar, 50 μm.

### Figure S13. The other two independent sets of results for Figure1-Figure5

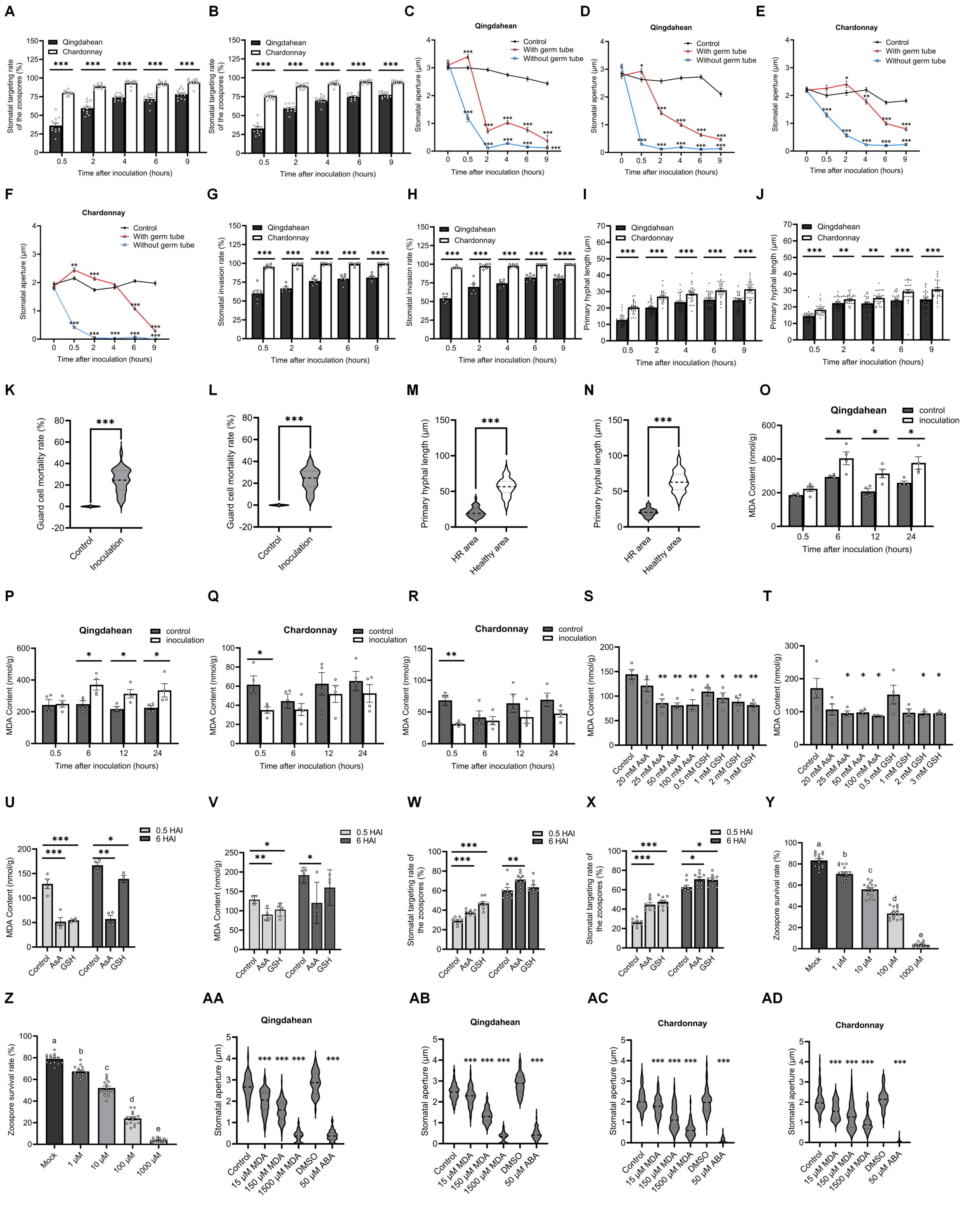


**(A-B)** Additional biological replicates for **(Figure 1)**; **(C-F)** Additional biological replicates for **(Figure 2)**; **(G-J)** Additional biological replicates for **(Figure 3)**; **(K-N)** Additional biological replicates for **(Figure 4)**; **(O-AD)** Additional biological replicates for **(Figure 5)**.
